# DNA methylation status classifies pleural mesothelioma cells according to their immune profile: implication for precision epigenetic therapy

**DOI:** 10.1101/2024.08.08.607174

**Authors:** Maria Fortunata Lofiego, Rossella Tufano, Emma Bello, Laura Solmonese, Francesco Marzani, Francesca Piazzini, Fabrizio Celesti, Francesca Pia Caruso, Teresa Maria Rosaria Noviello, Roberta Mortarini, Andrea Anichini, Michele Ceccarelli, Luana Calabrò, Michele Maio, Sandra Coral, Anna Maria Di Giacomo, Alessia Covre, the EPigenetic Immune-oncology Consortium Airc (EPICA) investigators

## Abstract

**Background:** co-targeting of immune checkpoint inhibitors (ICI) CTLA-4 and PD-1 has recently become the new first-line standard of care therapy of pleural mesothelioma (PM) patients, with a significant improvement of overall survival over conventional chemotherapy. The analysis by tumor histotype demonstrated a greater efficacy of ICI therapy in non-epithelioid (non-E) *vs* epithelioid (E) PM; although some E PM patients also benefit from treatment. This evidence suggests that molecular tumor features, beyond histotype, could be relevant to improve the efficacy of ICI therapy in PM. Among these, tumor DNA methylation emerges as a promising factor to explore, due to its potential role in driving the immune phenotype of cancer cells. Thus, we utilized a panel of cultured PM cells of different histotype, to provide preclinical evidence supporting the role of the tumor methylation landscape and of its pharmacologic modulation, to prospectively improve the efficacy of ICI therapy of PM patients.

**Methods:** the methylome profile (EPIC array) of distinct E (#5) and non-E (#9) PM cell lines was analyzed, followed by integrated analysis with their associated transcriptomic profile (Clariom S array), before and after *in vitro* treatment with the DNA hypomethylating agent (DHA) guadecitabine. The most variable methylated probes were selected to calculate the methylation score (CIMP index) for each cell line at baseline. Genes that were differentially expressed and methylated were then selected for gene ontology analysis.

**Results:** the CIMP index stratified PM cell lines in two distinct classes, CIMP (hyper-methylated; #7) and LOW (hypo-methylated; #7), regardless of their E or non-E histotype. Integrated analyses of methylome and transcriptome data revealed that CIMP PM cells had a substantial number of hyper-methylated, silenced genes, which negatively impacted their immune phenotype compared to LOW PM cells.

Treatment with DHA reverted the methylation-driven immune-compromised profile of CIMP PM cells and enhanced the constitutive immune-favorable profile of LOW PM cells.

**Conclusion:** the study highlighted the relevance of DNA methylation in shaping the constitutive immune classification of PM cells, that is independent from their histological subtypes. The identified role of DHA in shifting the phenotype of PM cells towards an immune-favorable state supports its role in clinical trials of precision epigenetic therapy combined with ICI.

## Introduction

Pleural mesothelioma (PM) is an aggressive cancer with a poor prognosis, characterized by a median survival rate of less than a year (1). Existing treatments, including chemotherapy, radiotherapy, and surgery, have shown limited effectiveness. However, the rising incidence of PM underscores the urgent need for novel and more effective therapies (2). Immune checkpoint inhibitors (ICI) have shown promise in treating various solid malignancies and are being explored for PM. Initial clinical results demonstrated that ICIs as monotherapies had limited success in improving overall survival (OS) of PM patients (3–5). However, a dual ICI approach combining the anti-cytotoxic T lymphocyte antigen-4 (CTLA)-4 monoclonal antibody (mAb) ipilimumab and the anti-programmed cell death (PD)-1 mAb nivolumab, tested in the phase III CheckMate 743 trial (NCT02899299), demonstrated greater efficacy, becoming the first-line therapy for unresectable chemo-naïve PM patients in several countries (6). In fact, combinatorial ICI regimens significantly improved the median OS of PM patients, compared to standard chemotherapy (18.1 *vs* 14.1 months). In particular, an improved survival benefit with combined ICI therapy *vs* chemotherapy was observed in PM patients with non-epithelioid (non-E) histology (18.1 *vs* 8.8 months), compared to those with epithelioid (E) histology (18.2 *vs* 16.7 months) (7). Despite these promising outcomes, the majority of PM patients failed to derive clinical benefit from ICI, or eventually develop resistance to ICI therapy. The different therapeutic effects of ICI might be attributed to an inability of treatments to reverse an “immune-cold” tumour microenvironment (TME), associated in PM with M2-macrophages and myeloid derived suppressor cells (MDSC) infiltration, (8), and lack of infiltrating effector T cells (9). A favorable influence of the immune contexture in PM clinical outcome is supported by different lines of evidence. First, low abundance of T-helper 2 and high cytotoxic T cells infiltration has been shown to promote improved OS in PM patients (10). In addition, a high number of B lymphocytes in TME and the presence of Tertiary lymphoid structures (TLS) predicts longer survival in E-PM patients (11). However, at present, factors determining PM response to immunotherapy remain elusive, but detailed response correlations with genome, transcriptome, methylome and immune landscape features are emerging. In this context, a quantitative molecular characterization of PM heterogeneity, by a deconvolution approach, identified this cancer as a mixture of E-PM- and non-E-PM-like cell population, whose proportions are highly associated with the prognosis (12). That study indicated that the molecular gradients represented by sarcomatoid (S)-score and epitheliod (E)-score were associated with distinct immune infiltration patterns in PM tumors, with the S-score linked to adaptive immune responses and exhausted T cell infiltration, which may have implications for the potential effectiveness of ICI therapy (12). In addition, the CpG island methylator phenotype (CIMP) index showed a positive correlation with the S-score, indicating the potential role of epigenetics as key regulator of the heterogeneity of PM tumors and immune microenvironment (8).

In this scenario, several studies highlighted that epigenetic drugs, mainly DNA hypomethylating agents (DHA), represented a strategy to increase tumor immunogenicity and reprogram the immune TME (13). In fact, it has been firmly demonstrated that epigenetic remodeling of cancer cells of different histotypes by DHA, such as decitabine and guadecitabine, induced/up regulated the expression of different immune molecules, resulting in improved immune recognition of tumor cells (14–17).

Based on this evidence, in this study we carried out a methylation profiling of human PM cell lines of different histological subtypes to develop an epigenetic classification based on the CIMP index. This approach identified two distinct CIMP (hyper-methylated) and LOW (hypo-methylated) PM classes, independent of their histological subtypes. We also found that the transcriptome was associated with the methylome-defined classes, revealing biological processes (BP) specifically activated or inhibited in each methylation-based subset. The most striking differences in CIMP *vs* LOW PM cell lines reflected the predominance of a negative regulation of immune activation in the former compared to the latter. In detail, an enrichment analysis of genes down-regulated by hyper-methylation in CIMP *vs* LOW PM cells, identified the inhibition of the antigen processing and presentation and B cell receptor signaling canonical pathways (CP), that could underpin an ICI-resistant phenotype in CIMP PM cells. Interestingly, we found that this immune compromised hyper-methylated profile in CIMP PM cell lines could be reverted by DHA treatment leading to activation of antigen processing and presentation pathways, and of inflammatory mediators. Noteworthy, the positive immunomodulatory effect of DHA was observed also in LOW PM cell lines, mainly associated with activation of Interferon-Mediated Signaling and antigen processing and presentation pathways.

Comprehensively, our results provide proof of principle evidence in favor of a methylation-based classification of PM cells as a novel approach to select/identify the PM tumors potentially more susceptible to ICI therapy. Moreover, the immunomodulatory activities of DHA treatment could represent a strategy to shift both CIMP and LOW PM cells towards an ideally more ICI-responsive phenotype, providing the rationale to explore a novel epigenetic-based ICI for treating PM patients, irrespective of their histology.

## Materials and methods

### Tumor cell lines

PM cell lines were established (#11) from pleural effusions of PM patients treated at the University Hospital of Siena, under approval by the Committee on Human Research or commercially purchased (#3) from American Type Culture Collection Cell Bank (ATCC, Rockville, MD, USA) and categorized into epithelioid (E; #5) and non-epithelioid (non-E; #9) histological subtypes. Cells were cultured using HAM’s F-12 medium (Euroclone, Milano), except for Meso 10, Meso 14, Meso 15 and Meso 16 that were grown in RPMI 1640 medium (Thermo Scientific, MA, USA). Both these mediums were supplemented with 10% heat-inactivated fetal bovine serum (FBS, Euroclone, Milan, Italy), 2 mML-glutamine (Thermo Scientific, MA, USA), and 100 µg/µL of penicillin/streptomycin (Euroclone, Milan, Italy). Cells were incubated at 37°C and 5 % CO2, they were passed when reached 80-90 % of confluency.

### *In vitro* epigenetic drug treatment

Treatment with guadecitabine was performed as previously described (18). Briefly, PM cell lines, tested for the absence of mycoplasma contamination (Roche, Indianapolis, USA), were seeded at 1×10^6^/mL in T75 cm^2^ tissue culture flasks (day 0) treated with 1 µM of guadecitabine (MedChemExpress) every 12 hours twice a day (day 1, day 2), and harvested at day 5 for all subsequent analyses. Control PM cell lines were maintained under similar experimental conditions without administering the drug.

### Clariom™ S human assay for gene-level expression profiling

Microarray profiling was performed on 500 ng of DNAase I digested RNA using the Affymetrix Clariom™ S human microarray platform (Affymetrix, Santa Clara, CA). Each biotin-labeled sense target was hybridized onto a single GeneChip® Clariom™ S Affymetrix human microarray. After hybridization, the microarrays underwent washing steps to remove any nonspecifically bound material and then subjected to image acquisition using the Affymetrix Fluidics Station 450. The acquired images were processed using the Affymetrix GeneChip® Command Console software (provided by Thermo Fisher Scientific, Inc).

### Genome-wide methylation analysis screening: the Infinium MethylationEPIC array

Genome-wide methylation analysis was performed on 500 ng of DNA isolated from #14 human PM cell lines of different histotype using QIAamp DNA Blood Mini Kit (Qiagen, Hilden, Germany) by the Infinium Human Methylation EPIC 850k (EPIC) array, according to the manufacturer’s standard protocols. In detail, EPIC 850k array is based on Illumina’s BeadChip technology, where dual-probe design captured the full spectrum of DNA methylation at the single-CpG-site level at high resolution. The Infinium design is based on specific probes interrogating each CpG site, the signal intensity, emitted by their interaction is then measured to generate beta values (β), defined as β = M/(M+U+100), where M is the intensity corresponding to methylated and U to unmethylated, representing the relative degree of methylation at a locus. β values have a value ranging from 0 to 1, representing fully unmethylated and methylated states, respectively. The β values were normalized by using the functional normalization approach (19). Then, we used the β values of cell lines (at the baseline) to estimate the most variable probes and calculated a methylation score by summing the methylation level of selected probes. Raw intensities were processed using minfi R package (v 1.42.0) (20). The methylation data were normalized by using the functional normalization approach implemented in the function preprocessFunnorm() of the R package minfi (20). Then, the probes with low quality, located in the X and Y chromosomes, known to have single nucleotide polymorphisms (SNPs) at the CpG site and to be cross-reactive were filtered out.

### Data analysis

#### Classification of PM cell lines based on the CIMP index

The normalized and filtered probes were used to estimate the most variable ones based on the difference between the 10th percentile and 90th percentile of the data. Almost 1% of the total probes were selected as the most variable and used to calculate the methylation score (CIMP-index) for each cell line at baseline by summing the β values across these selected probes. The PM cell lines were then categorized into two groups according to the median CIMP index: those with a methylation score above the median (CIMP) and those with a methylation score below the median (LOW).

#### Differential methylation analysis

Differential methylation analysis was performed to identify differentially methylated (DM) probes between CIMP *vs* LOW untreated PM cell lines, as well as between DHA-treated and untreated PM cell lines within both CIMP and LOW groups, using the R package limma (21) by fitting a linear model to the matrix of M values. Probes with a *p*-value<0.05 and |log2FC| >= 1 were selected as DM. Gene ontology (GO) enrichment analysis was performed using the gometh() function of the R package missMethyl (22) focusing on DM probes overlapping the promoter region (5’UTR, TSS1500, TSS200). Biological processes (BP) with a *p*-value<0.01 were then selected.

#### Differential expression analysis

Differential expression analysis was performed to identify differentially expressed genes between CIMP *vs* LOW untreated PM cell lines, as well as between DHA-treated and untreated PM cell lines within both CIMP and LOW groups, using Transcriptome Analysis Console (TAC) Software (Applied Biosystems, Thermo Fisher Scientific). Genes with a *p*-value<0.05 and |log2FC| >= 1 were selected as differentially expressed.

#### Integrative analysis of methylation and expression data

Results of the methylation and expression analyses for each comparison were integrated by selecting genes that were both differentially expressed and differentially methylated at the promoter region. Starburst plots displayed DM probes overlapping the promoter region. The -log10(*p*-value) of the up-regulated genes and the log10(*p*-value) of down-regulated genes were plotted against the -log10(*p*-value) of the hyper-methylated probes and the log10(*p*-values) of down-methylated probes. These modules of expression/methylation concordant genes were functionally analyzed using Ingenuity Pathway Analysis tool (IPA 8.5, www.ingenuity.com).

#### GO terms analysis

IPA Core analysis was performed to identify, at transcriptomic level, upstream regulators (UR) activated (Z-score[≥2) or inhibited (Z-score[≤−2) at the constitutive level in CIMP *vs* LOW PM cell lines, as well as in both CIMP and LOW classes before and after DHA treatment. Enrichment of GO terms, considering BP, was conducted utilizing the EnrichR web-tool (23). Significative BP (*p*-value<0.05) were ranked based on their combined score value, which was calculated by multiplying the log of the *p*-value (computed with the Fisher exact test) by the Z-score from our correction to the test. The top 50 BP were then analyzed. Moreover, IPA Core analysis was performed on hyper-methylated down-regulated genes, identified by integrative analysis, to define CP inhibited (Z-score[≤−2) at the constitutive level in CIMP *vs* LOW PM cell lines and on hypo-methylated up-regulated genes, identified by integrative analysis, to define CP activated (Z-score[≥2) in CIMP and LOW PM cell lines after DHA treatment.

#### Analysis in PM lesions from the TCGA-MESO cohort

Illumina 450K methylation array data of The Cancer Genome Atlas (TCGA) MESO cohort (87 samples) were downloaded with the TCGABiolinks R package (24). Then, the probes located in the X and Y chromosomes and in the open sea regions, with SNPs at the CpG site and to be cross-reactive, and bowtie2 multi-mapped were removed. The filtered probes were used to estimate the most variable ones based on the difference between the 10th percentile and the 90th percentile of the data. 2285 sites were selected as the most variable sites and used to calculate the methylation score for each sample. The methylation score was calculated by summing the β values across the variable sites for each sample. The top 25% and the bottom 25% of samples (22 samples for each group), according to the methylation score, were selected respectively as hyper-methylated (CIMP) and hypo-methylated samples (LOW).

The expression data of the TCGA-MESO cohort were downloaded from cBioPortal (http://www.cbioportal.org, accessed on 01 August 2024). The file ‘data_mrna_seq_v2_rsem.txt’ was downloaded and used to explore the differences in the expression profile between the CIMP and LOW samples and to estimate the abundance of eight immune and two stromal cell populations in the tissues.

The differential expression analysis was performed by applying the Wilcoxon Test and comparing the CIMP samples to the LOW ones. Then, the genes sorted according to the Wilcoxon statistics were used for a gene set enrichment analysis with the clusterProfiler R package (25). The abundance of the cell populations in the tissue of each sample was estimated with the MCPcounter R package deconvolution analysis (26). The Student’s t-test was performed to assess if there was a significant difference between the groups.

### Gene expression analysis by quantitative Real-Time RT-PCR

Relative quantification by Real-Time PCR was performed on 21.8 ng of retrotranscribed RNA in a final volume of 20 μl TaqMan Fast Advanced Master Mix (Applied Biosystems, CA, United States). Relative gene expression levels were quantified by the 2-ΔΔCT method, where ΔCT was the difference between CT values of the gene target and the reference gene β-actin; ΔΔCT represented the difference between ΔCT of treated *vs* untreated cells (FC). Gene expression was considered up-(FC ≥1.5) or down-regulated (FC ≤ 0.5) when its level exhibited a 1.5-fold increase

or decrease. All the relative quantification analyses were performed using the QuantStudio ™ 5 Real-Time RT-PCR System (Applied Biosystems ™, CA, USA) and its analysis software. Gene-specific probes for ERVs and ISGs were selected form Taqman Gene Expression Assay database (Applied Biosystem, Foster City, CA, USA) (**Table 1**) or designed by Custom Assay Design (Applied Biosystem, Foster City, CA, USA) (**Table 2**).

**Table 1.**
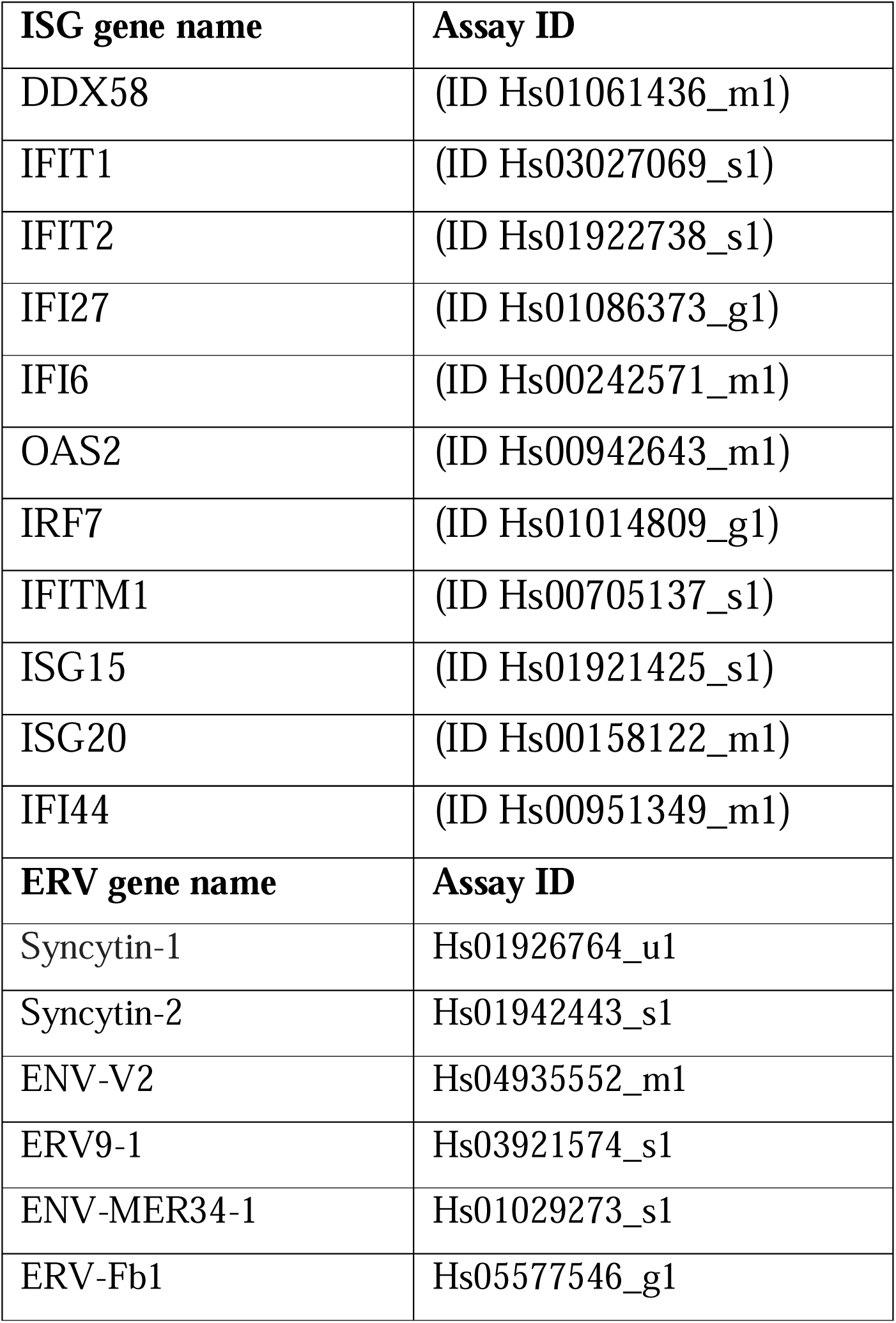
Gene specific assays for ISGs and ERVs expression.

**Table 2.**
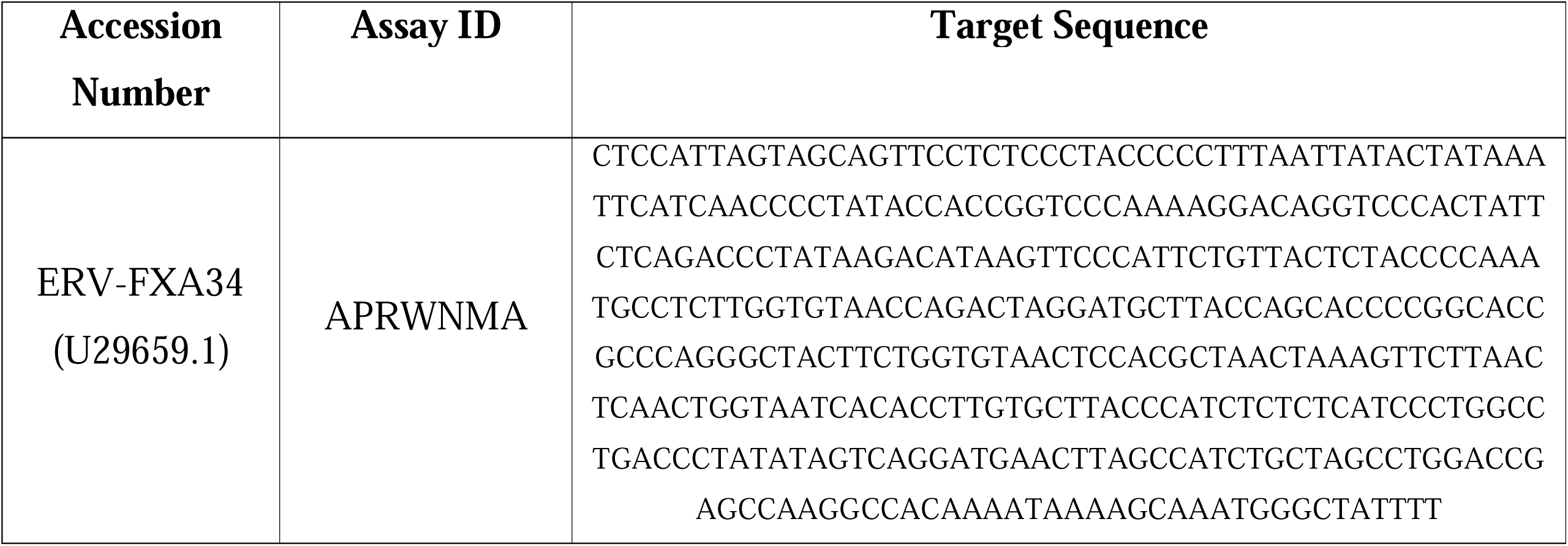
Customized gene specific assay for ERV-FXA34 expression.

## Results

### Methylation-based classification of PM cell lines

The methylome analysis identified #7,850 probes as the most variably methylated among all investigated PM cell lines, that were used to calculate the CIMP-index of each cell line. PM cell lines were then stratified into two classes: hyper-methylated (CIMP; #7) and hypo-methylated (LOW; #7) based on their CIMP-index higher or lower than median value, respectively (**Figure 1 A**). Principal component analysis (PCA) demonstrated that this methylation-based PM cell lines stratification was independent from their histopathological categorization, indeed E and non-E PM cell lines were homogeneously distributed among above median and below median CIMP index values (**Figure 1 B**). Among the mapped methylated probes, #63,154 were significantly (*p*-value<0.05) differentially methylated (DM) (#58,710 hyper- and #4,444 hypo-methylated) in CIMP *vs* LOW PM cell lines confirming the stratification of these cell lines into PCA-defined groups, independent of their histological classification into E and non-E subtypes (**Additional Figure 1 A**).

**Figure 1.**
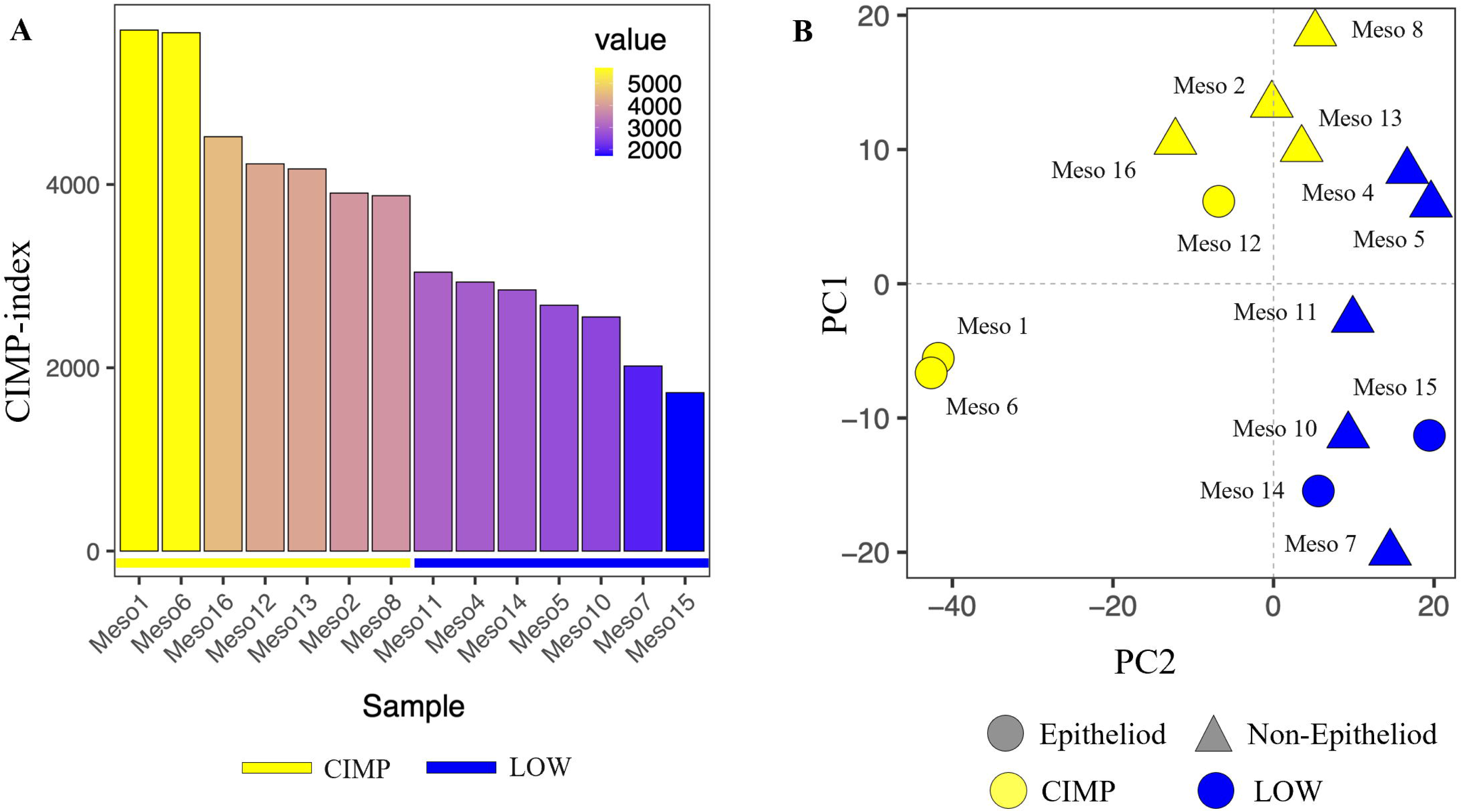
Analysis of methylation patterns (CIMP-index) in PM cell lines. Starting from EPIC array data, the interleaved bar plot represented the CIMP-index, for each cell line, calculated by summing the β values of the most variable methylated probes, that represented approximately 1% of the total probes. The β values were normalized by using the functional normalization approach. Each bar represented a single E or non-E PM cell line at constitutive level (**A**). Dimensionality reduction was performed applying PCA on the most variable methylated probes among all PM cell lines. Each symbol on the graph represented a cell line categorized by its methylation status: CIMP (yellow) and LOW (blue), and by its histopathological variant: E (circle) and non-E (triangle) (**B**).

To explore the biological effects of the identified DM probes, we explore their distribution across the genome according to their relation to CpG islands (**Additional Figure 2 A, B**) and gene functional region (**Additional Figure 2 C, D**) in CIMP *vs* LOW PM cell lines. Results showed that the hyper-methylated probes in the CIMP PM cell lines compared to the LOW ones were enriched in the 1stExon (*p*-value < 2.2e-16, proportion test), 5’ UTR (*p*-value< 2.2e-16, proportion test), TSS200 (*p*-value< 2.2e-16, proportion test) and TSS500 (*p*-value< 2.2e-16, proportion test) compared to the hypo-methylated ones. In addition, the hyper-methylated probes were mainly located in CpG island regions (*p*-value< 2.2e-16, proportion test) and north shore regions (*p*-value=7.474e-08, proportion test). To comprehensively investigate the methylation-driven biological differences of CIMP *vs* LOW PM cell lines, an enrichment analysis was conducted on DM probes located in the promoter region. Results highlighted multiple processes involved in cell-cell interaction signaling, cell differentiation and chemical synaptic transmission signaling, impacted by hyper-methylated genes, or on biosynthetic and transcription process impacted by hypo-methylated genes, in CIMP *vs* LOW PM cell lines (**Figure 2 A, C**). Focusing on the immune-related (IR) processes category, comparing CIMP *vs* LOW PM cell lines, the enriched processes impacted by hyper- or hypo-methylated probes were mainly involved in the regulation of humoral immune response, T cell selection and leukocyte migration/chemiotaxis or inflammatory response to antigenic stimulus, dendritic cell (DC) differentiation and production of molecular mediator of immune response, respectively (**Figure 2 B, D**).

**Figure 2.**
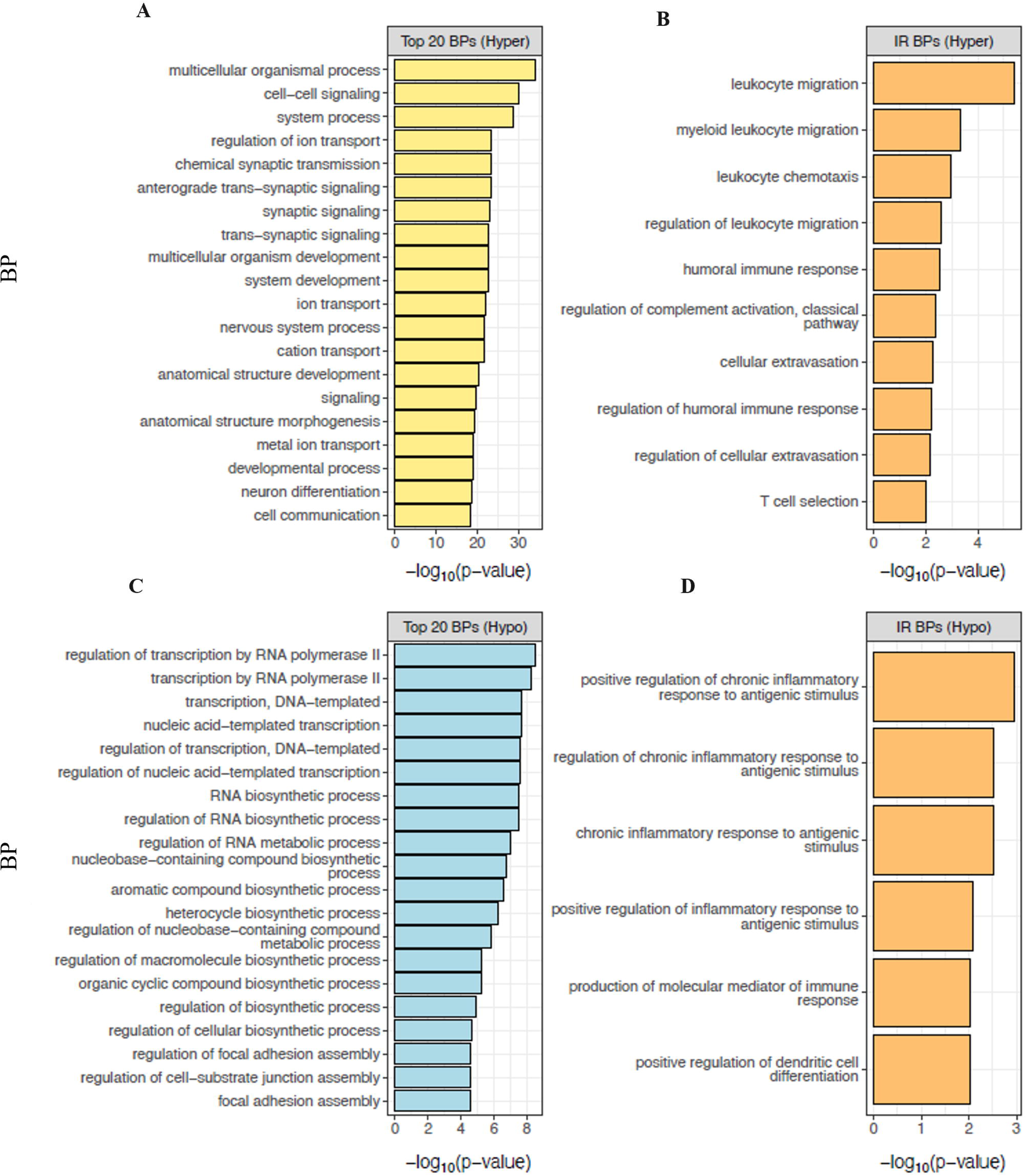
Enrichment analysis of hyper-methylated (A, B) and hypo-methylated (C, D) probes in the promoter region of CIMP *vs* LOW PM cell lines. The top 20 BP enriched by hyper-(**A)** or hypo- (**C**) methylated probes in CIMP *vs* LOW PM cell lines were identified. The top 10 IR-BPs enriched by hyper- methylated probes (**B**) and the top 6 IR-BP enriched by hypo-methylated (**D**) probes in CIMP *vs* LOW PM cell lines were identified

### Transcriptomic profiles of CIMP *vs* LOW PM cell lines

To comprehensively examine the phenotypic differences of DM PM cell lines, a comparative gene expression analysis of CIMP *vs* LOW PM cell lines was performed, identifying #649 up-regulated genes and #595 down-regulated genes. Bioinformatic IPA core analysis was carried out to predict the constitutive functional status (activation or inhibition) of UR in CIMP *vs* LOW PM cells lines and identify their associated BP (**Figure 3 A**). Results showed that in CIMP PM cells, activated (Z-score≥2) UR were mainly associated with the BP involved in shutting down of the immune response (i.e., negative response to type I interferon, negative regulation of T-helper (Th)-17 type immune response, and negative regulation of T cell cytokine production), in cancer cell invasion and drug resistance (i.e., Hippo signaling pathway), and in tumor progression (i.e., TORC1 signaling) (**Figure 3 A; Additional File 1:Table S1, S2**). Likewise, UR inhibited (Z-score≤−2) in CIMP PM cell lines enriched BP linked to the acute inflammatory response, suppression of Th-1 cells and secretion of pro-inflammatory cytokines (i.e., IL-6, Th-1 cytokine, monocyte chemotactic protein-1) (**Figure 3 B; Additional File 1: Table S3, S4**). Similarly, in CIMP *vs* LOW PM cell lines, several CP were inhibited, including IL-17 signaling, PD-1/PD-L1 cancer immunotherapy pathways, T Cell Receptor and B Cell Receptor signaling, that exert a pivotal role in coordinating an efficient adaptive cell-mediated immune response, and non-canonical NF-κB signaling, implicated in T cell development. Additionally, the mitotic G2/M phases CP were also inhibited (**Additional File 2: Table S1**). Among the predicted activated CP in CIMP *vs* LOW PM cell lines were those related to RORA, RAS processing and PPAR signaling, primarily involved in tumor cell proliferation and invasion (22, 23). Notably, oncogenic NOTCH1 signaling and cellular stress response pathways (i.e., cytoprotection by HMOX1, and p75 NTR receptor-mediated signaling) were activated in CIMP compared to LOW PM cell lines (**Additional File 2: Table S1**).

**Figure 3.**
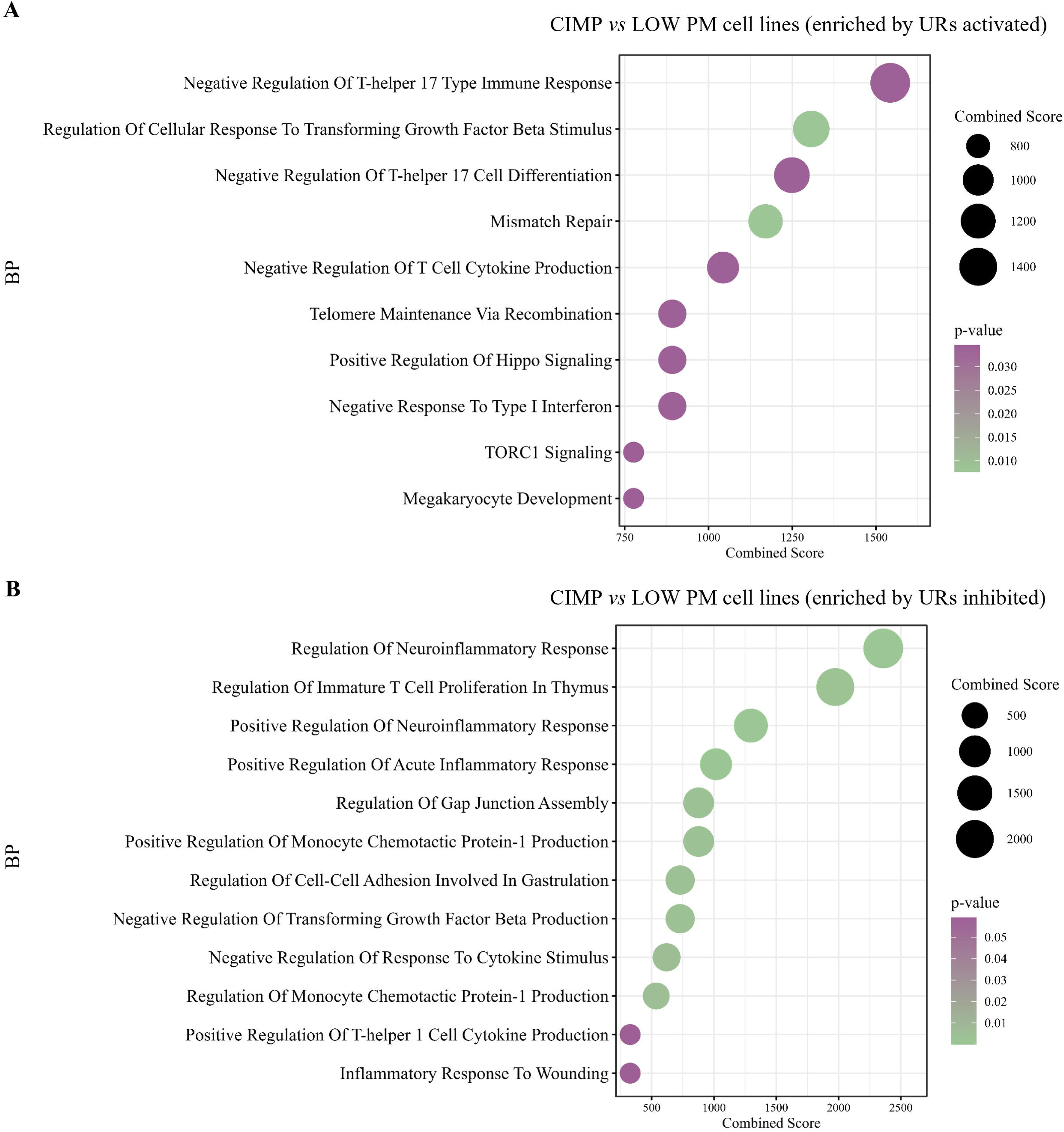
Representative functional categories enriched by differentially activated or inhibited UR in CIMP *vs* LOW PM cell lines. Dot plot of representative GO terms enriched by activated (**A**) or inhibited (**B**) UR in CIMP *vs* LOW PM cell lines. The color scale from purple to green indicated the ranges of significant *p*-value (p<0.05) and the size of each single dot correlated with the combined score.

To investigate the role of the LOW and CIMP methylator phenotype and of its related immune profile identified in PM cell lines, in contributing to shape the methylation phenotype and the immune contexture of PM tumors, the multi-omics profiling of #87 tumor lesions of PM patients from the TCGA-MESO cohort was exploited. Patients were stratified according to their tumor methylation score, and classified in hyper-methylated (CIMP; #22, top 25%) and hypo-methylated (LOW; #22, bottom 25%) classes. This classification was independent from the PM histological subtypes. The opposite immune-low and immune-high transcriptional profile identified in CIMP and LOW PM cell lines, respectively, was recapitulated *ex vivo* in CIMP and LOW PM tumors (**Figure 4 A)**. Indeed, genes expressed in the LOW PM lesions enriched pathways associated with immune response, whereas the CIMP lesions were characterized by enhanced proliferation (**Figure 4 A**). In addition, a significantly (*p*<0.05) higher infiltration of cytotoxic lymphocytes, together with B, myeloid DC, and endothelial cells, was identified by TME deconvolution in LOW compared to CIMP PM lesions (**Figure 4 B**).

**Figure 4:**
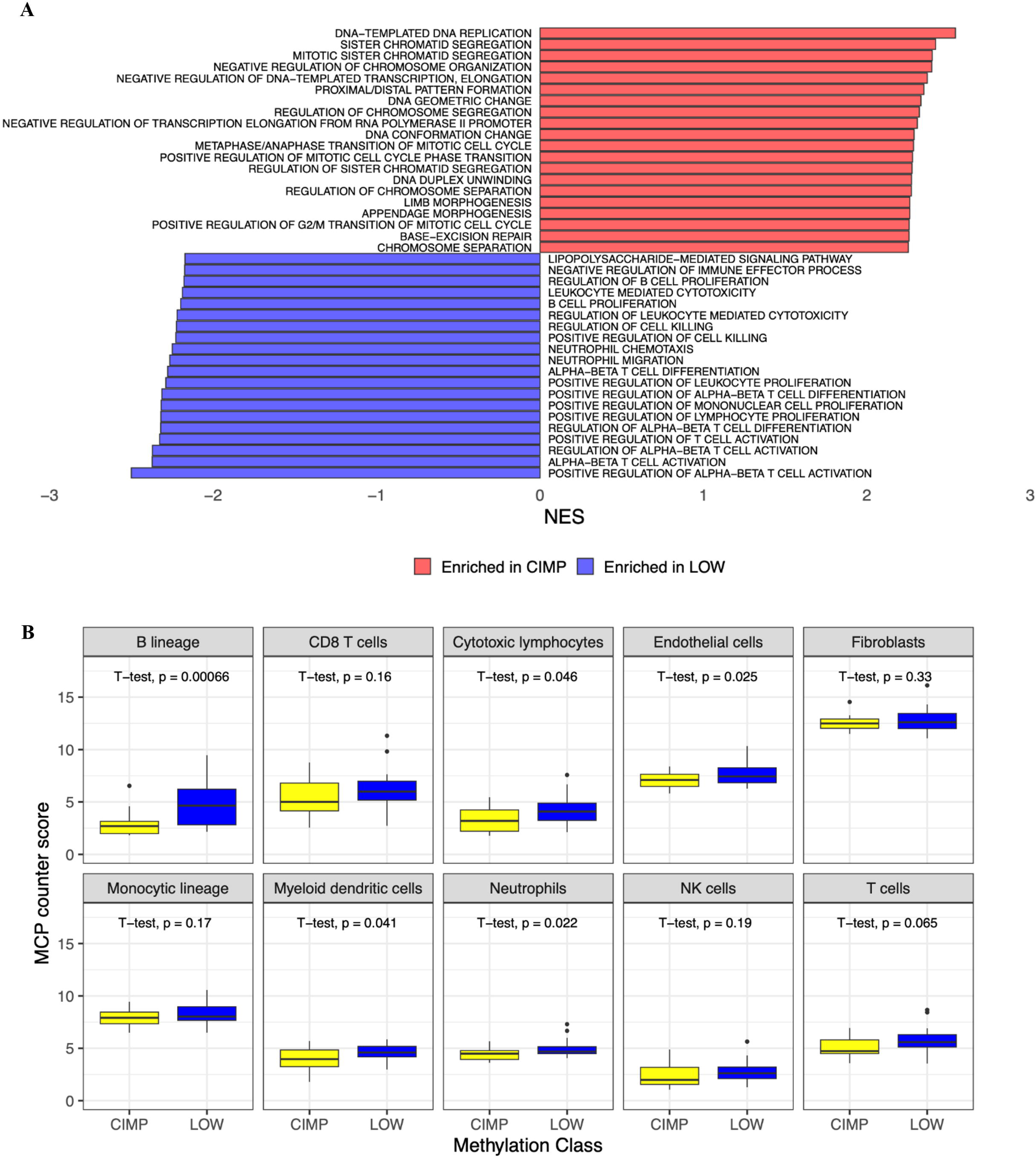
Immune phenotypic characterization of methylation-classified tumors from the TCGA-MESO cohort. Gene set enrichment analysis of the differentially expressed genes in the top 25% hyper-methylated (CIMP) *vs* the bottom 25% hypo-methylated (LOW) tumor samples from PM patients of the TCGA-MESO cohort. The top 20 BP were shown (**A**). Tumor microenvironment deconvolution of immune and stromal cell fractions in CIMP and LOW PM tissues from the TCGA-MESO cohort’ patients (**B**).

### Integrative analysis of promoter methylation level and transcriptomic profiles in CIMP *vs* LOW PM cell lines

Integrative analyses of methylome and transcriptomic data in CIMP *vs* LOW PM cell lines were performed to study the direct involvement of promoter methylation in the poorly immunogenic profile observed in CIMP PM cell lines (**Figure 5 A**). Particularly, an enrichment analysis was conducted on genes (#217) whose expression was directly down-regulated by hypermethylation (**Figure 5B**) in CIMP *vs* LOW PM cell lines, identifying inhibited (Z-score ≤ -2) CP mainly associated with class I MHC-mediated antigen processing and presentation, IL-17 signaling, DC maturation and BCR signaling (**Additional File 3: Table S1)**. Accordingly, the enrichment analysis in CIMP *vs* LOW PM cells, identified inhibited (Z-score ≤ -2) UR with a predominant influence on tumor necrosis factor-mediated signaling pathway, type II IFN-mediated signaling, regulation of cytokine production involved in inflammatory response, and myeloid DC differentiation BP (**Additional File 3: Table S2, S3**). Moreover, invasive/proliferative processes and extracellular matrix disassembly were also inhibited by hyper-methylated silenced genes (**Figure 5 B, Additional File 3: Table S3**).

**Figure 5.**
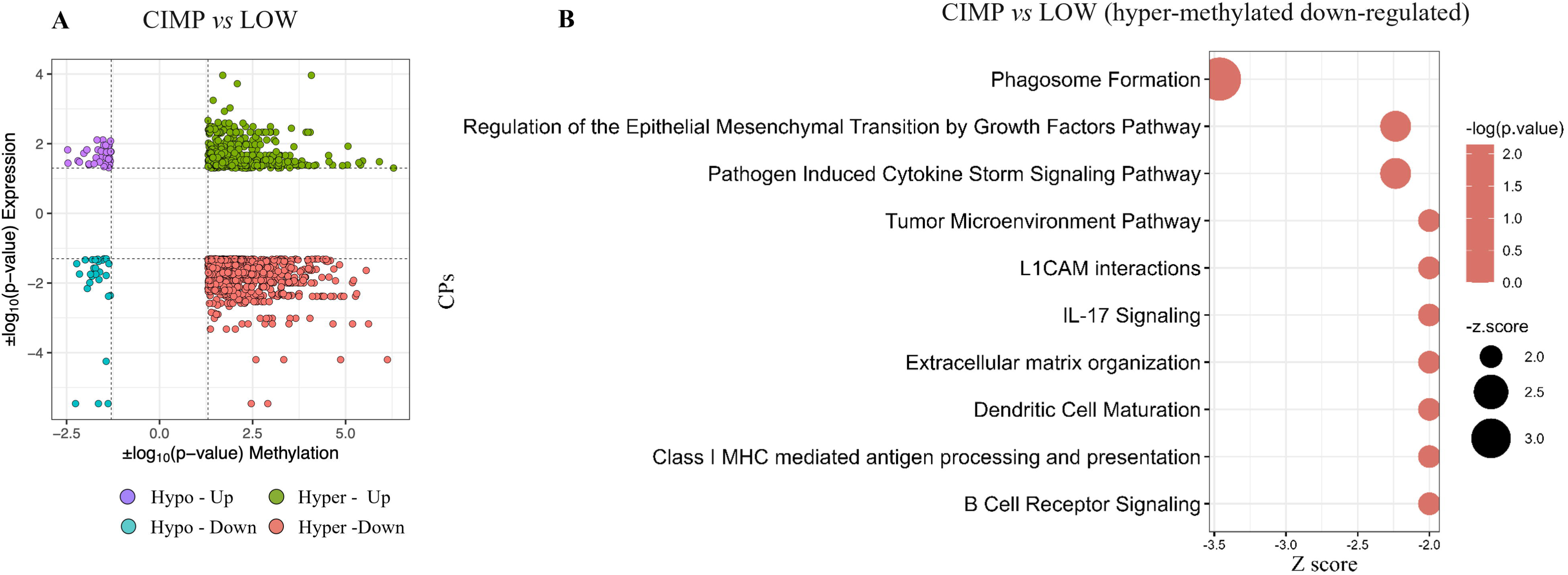
Integrative methylation and expression analysis. The starburst plot showed results of the integrative analysis of promoter methylation and transcriptomic data in CIMP *vs* LOW PM cell lines. x and y axis represented the |log_10_(*p*-value)| of methylation and the |log_10_(*p*-value)| of expression (**A**), respectively. CP enriched by hyper-methylated and down-regulated genes in CIMP *vs* LOW PM cell lines were identified by IPA core analysis (**B**).

### Remodeling of methylation profile in CIMP and LOW PM cell lines by DHA treatment

To test whether the hyper-methylated profile of CIMP PM lines could be reverted, we compared the methylation profiles of each CIMP and LOW PM cell line, before and after treatment with guadecitabine. Results demonstrated a more frequent reduction of global methylation in CIMP compared to LOW PM cell lines following DHA treatment (**Figure 6 A**); indeed, among the significantly (*p*<0.05) DM probes, a total of #152,420 and #5,765 probes were hypo-methylated in DHA-treated CIMP and LOW PM cell lines, respectively. PCA stratification analysis revealed that both CIMP and LOW PM cell lines clustered according to DHA treatment, regardless of their histopathological classification (**Figure 6 B, C**).

**Figure 6.**
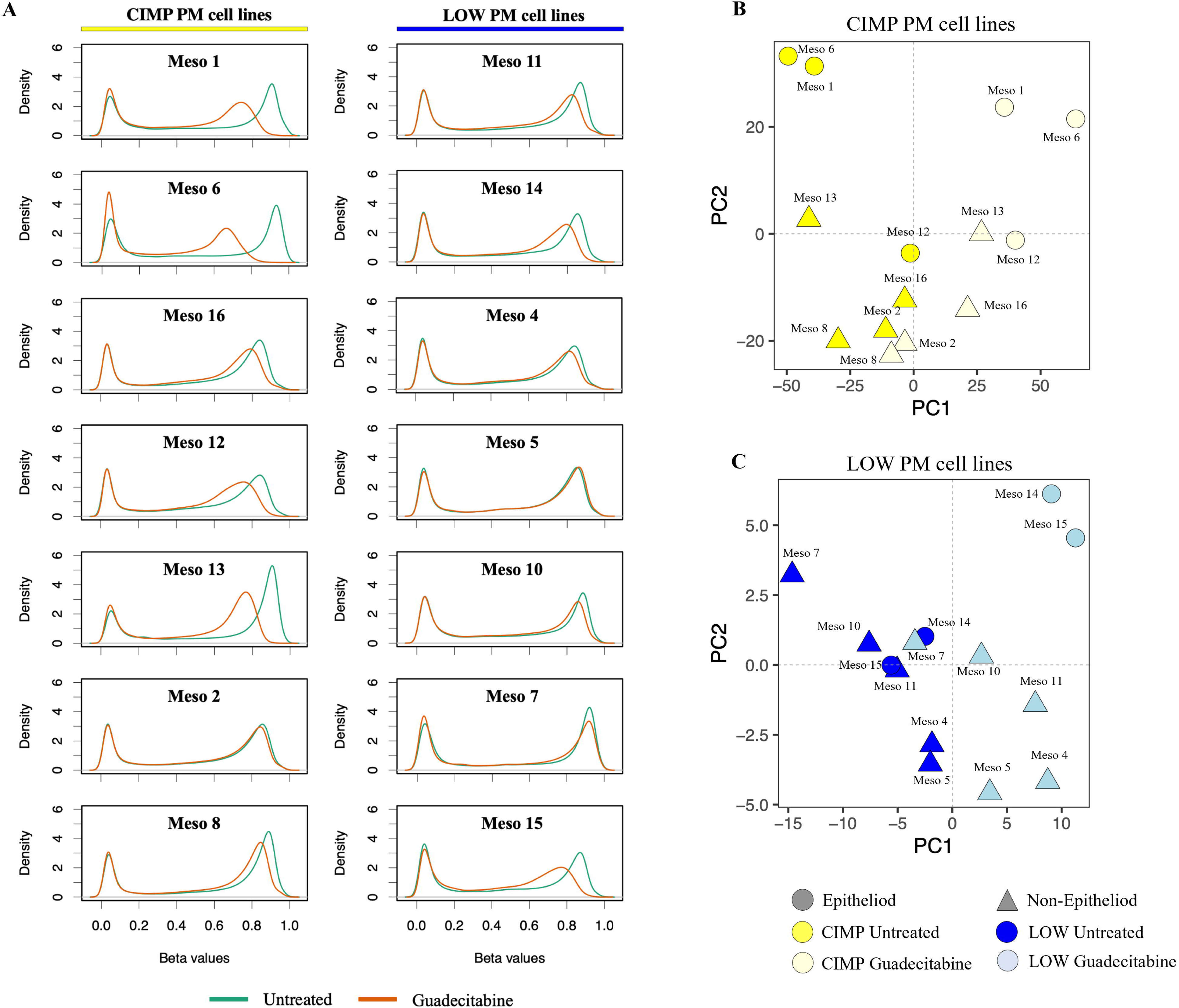
Modulation of methylation in DHA-treated *vs* untreated PM cell lines. Genomic DNA was extracted from untreated and DHA-treated #14 PM cell lines and used to investigate the methylation profile through Infinium Methylation EPIC array. The density of methylation β value for each CIMP and LOW cell line both at the baseline (green) and after guadecitabine treatment (orange) was plotted (**A**). Dimensionality reduction was performed by applying PCA on the DM probes among DHA-treated and untreated CIMP (**B**) and LOW (**C**) PM cell lines. E-PM (circle) or non-E-PM (triangle).

An enrichment analysis was performed on DM probes in DHA-treated *vs* untreated CIMP and LOW PM cell lines to investigate the BP associated with demethylation. The enrichment analysis of hypo-methylated probes in the promoter region of DHA-treated CIMP PM cells showed an impact on cellular developmental and reproductive/sexual processes as well as multicellular organismal processes (**Figure 7 A**), and an enrichment of type 2 immune response, leukocyte degranulation, mast cell mediated immunity and myeloid leukocyte differentiation (**Figure 7 B**). The same analysis of hypo-methylated probes of DHA-treated LOW PM cells revealed an impact on chromatin organization, cellular component assembly involved in morphogenesis, lymphocytes co-stimulation and phosphatidylinositol-3-phosphate biosynthetic process (**Figure 7 C**), as well as multiple IR-categories, including T cell co-stimulation, regulation of NK cell-mediated cytotoxicity, and regulation of T cell extravasation (**Figure 7 D**).

**Figure 7.**
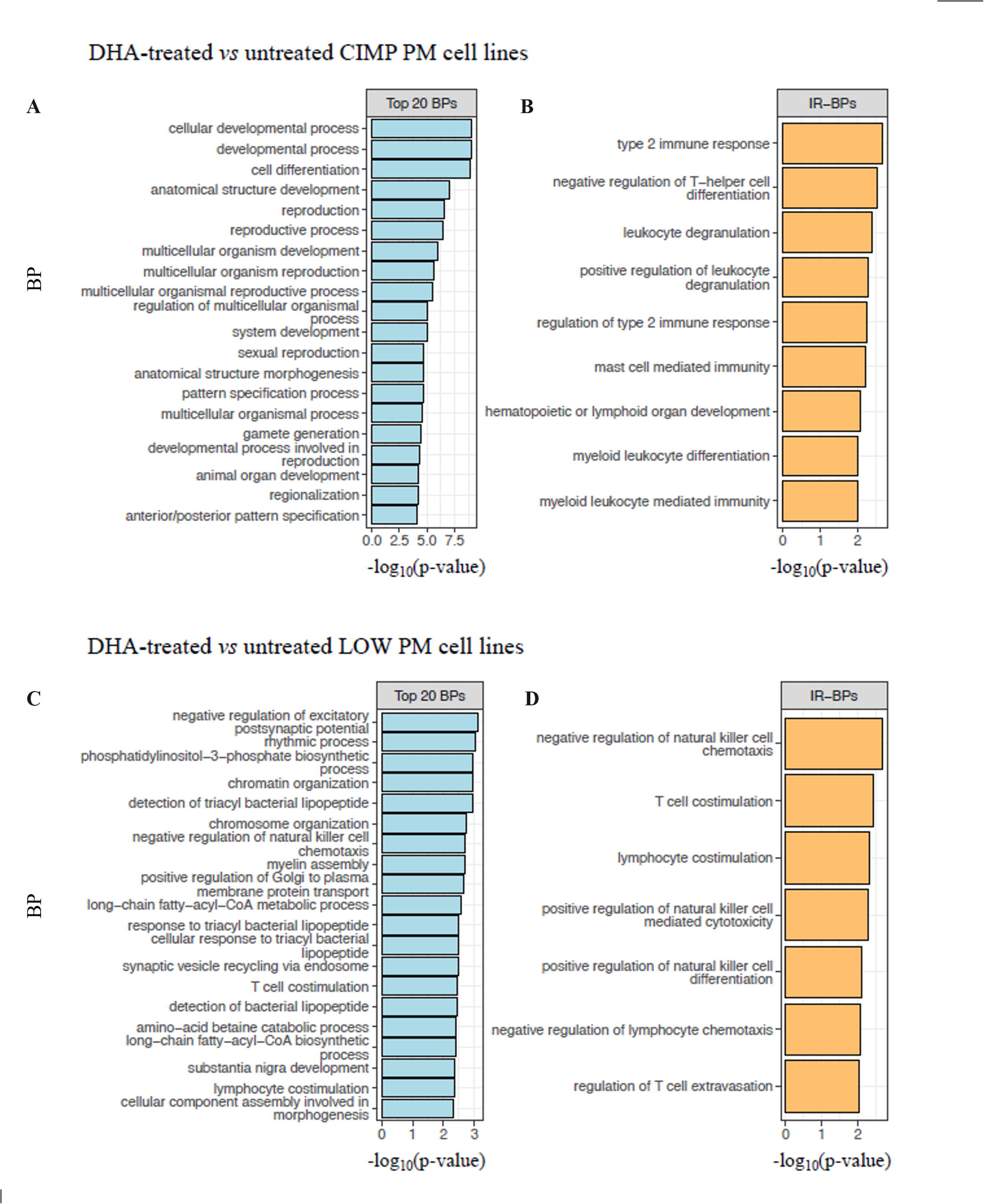
Enrichment analysis of hypo-methylated probes in the promoter region of DHA-treated *vs* untreated CIMP and LOW PM cell lines. The top 20 BP enriched by hypo-methylated probes of DHA-treated *vs* untreated genes in CIMP (**A**) and LOW (**C**) PM cell lines. The top 9 IR-BPs enriched by hypo-methylated probes of DHA-treated *vs* untreated genes in CIMP PM cell lines (**B**), and the top 7 IR-BP enriched by hypo-methylated probes of DHA-treated *vs* untreated genes in LOW PM cell lines (**D**).

### Transcriptomic profiles of CIMP and LOW PM cell lines after DHA treatment

Transcriptomic analysis was performed to comprehensively explore the effects of guadecitabine on the gene expression profile of CIMP and LOW PM cell lines. Results showed that, among #21,448 investigated genes, #2,358 and #3,034 were significantly (*p*<0.05) modulated by guadecitabine in CIMP and LOW PM cell lines, respectively. In detail, #1,147 and #1,211 DEGs were up- and down-regulated, respectively, in CIMP PM cell lines, while #1,431 and #1,603 DEGs were up- and down-regulated, respectively, in LOW PM cell lines. It is noteworthy that in the CIMP PM cells, which consistently displayed a strong immune suppression signature, DHA treatment primarily enhanced the activation of CP related to MHC class I and class II antigen processing and presentation, as well as the cGAS-STING signaling pathway, which is crucial for IFN production and host antiviral responses. (**Additional File 4: Table S1**). For this purpose, guadecitabine activated specifically IFN alpha, beta and gamma signaling, beyond the CD28 costimulatory pathway, TCR and NK cell signaling (**Additional File 4: Table S1**). In addition, BP enriched by activated UR were mainly involved in chemokine production, cytokine-mediated signaling and positive regulation of acute inflammatory response (**Figure 8A; Additional File 4: Table S2, S3**). Moreover, only 4 CP were inhibited by guadecitabine in CIMP PM cells, mainly affecting cell invasion and proliferation processes (i.e., Wnt/β-catenin pathway and PPARα signaling) (**Additional File 4: Table S1)**, and inhibited UR enriched for BP mainly involved in biosynthetic, metabolic processes and in the regulation of TGF-β receptor signaling (**Additional File 4: Table S4, S5)**. Contextually, DHA treatment also shaped the immune-favorable constitutive profile of LOW PM cell lines. In detail, CP activated by DHA affected a large set of IR-pathways (i.e., TCR signaling, cGAS-STING signaling pathway, DC maturation, MHC class I/II antigen presentation, NK Cell Signaling, ICOS-ICOSL Signaling in T-helper cells) (**Additional File 4: Table S6)** and besides these, IFN-mediated signaling pathway and cytokine production were enriched by DHA- activated UR (**Figure 8B, Additional File 4: Table S7, S8**). Analysis of CP and UR inhibited by DHA, in LOW PM cell lines, were mainly involved in metabolic and proliferative pathways (**Additional File 4: Table S6, S9-S10**).

**Figure 8.**
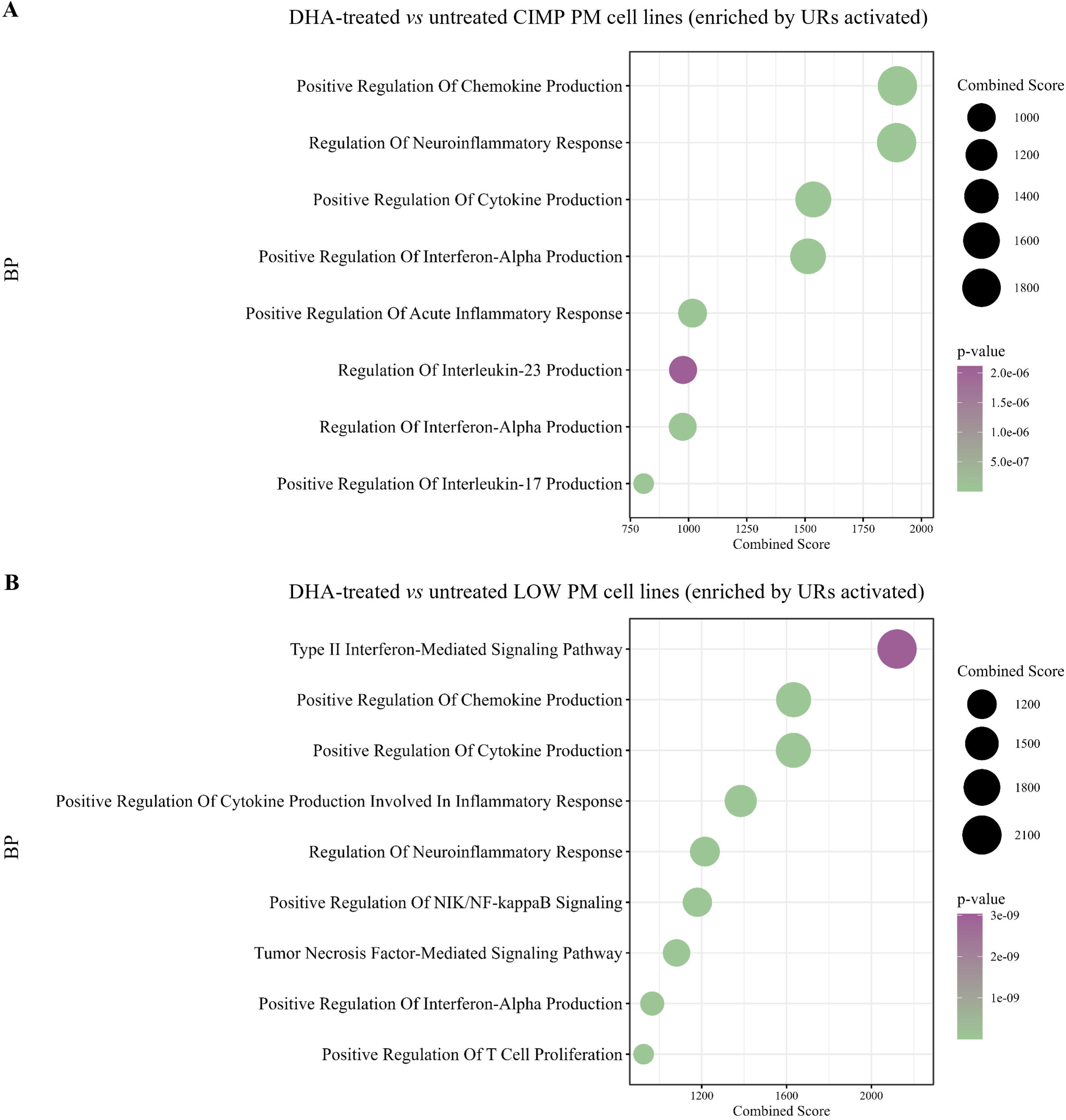
Transcriptomic landscape of CIMP and LOW PM cells after DHA treatment. Dot plots display the representative BP enriched in the GO BP signatures in CIMP (**A**) and LOW (**B**) PM cell lines after guadecitabine treatment, respectively. The color scale from purple to green indicates the ranges of significant *p*-value (*p*<0.05), and the size of each single dot correlates with the combined score.

One of the main effects exerted by DHA is the induction of the viral mimicry phenomenon through the up-regulation of human endogenous retroviruses (HERV), that activates an antiviral state producing type I and III IFNs and promoting the transcription of interferon-stimulated genes (ISG) (24,25). Consistent with the enrichment of BP related to IFN-mediated signaling pathway observed in both DHA-treated CIMP and LOW PM cell lines, a global up-regulation in the expression of HERV (**Figure 9 A**) and ISG (**Figure 9 B**) genes was observed across all PM cell lines, with a greater extent in LOW PM cell lines. In particular, although not statistically significant, a higher median fold change (mFC) was observed in treated *vs* untreated LOW compared to CIMP PM cell lines for HERV (mFC=4.44 *vs* mFC=2.39) and for ISGs (mFC=1.97 *vs* mFC=1.71), respectively. The modulation of HERV and ISG expression appeared independent of histological classification in both CIMP and LOW PM cell lines (**Figure 9 A, B**).

**Figure 9.**
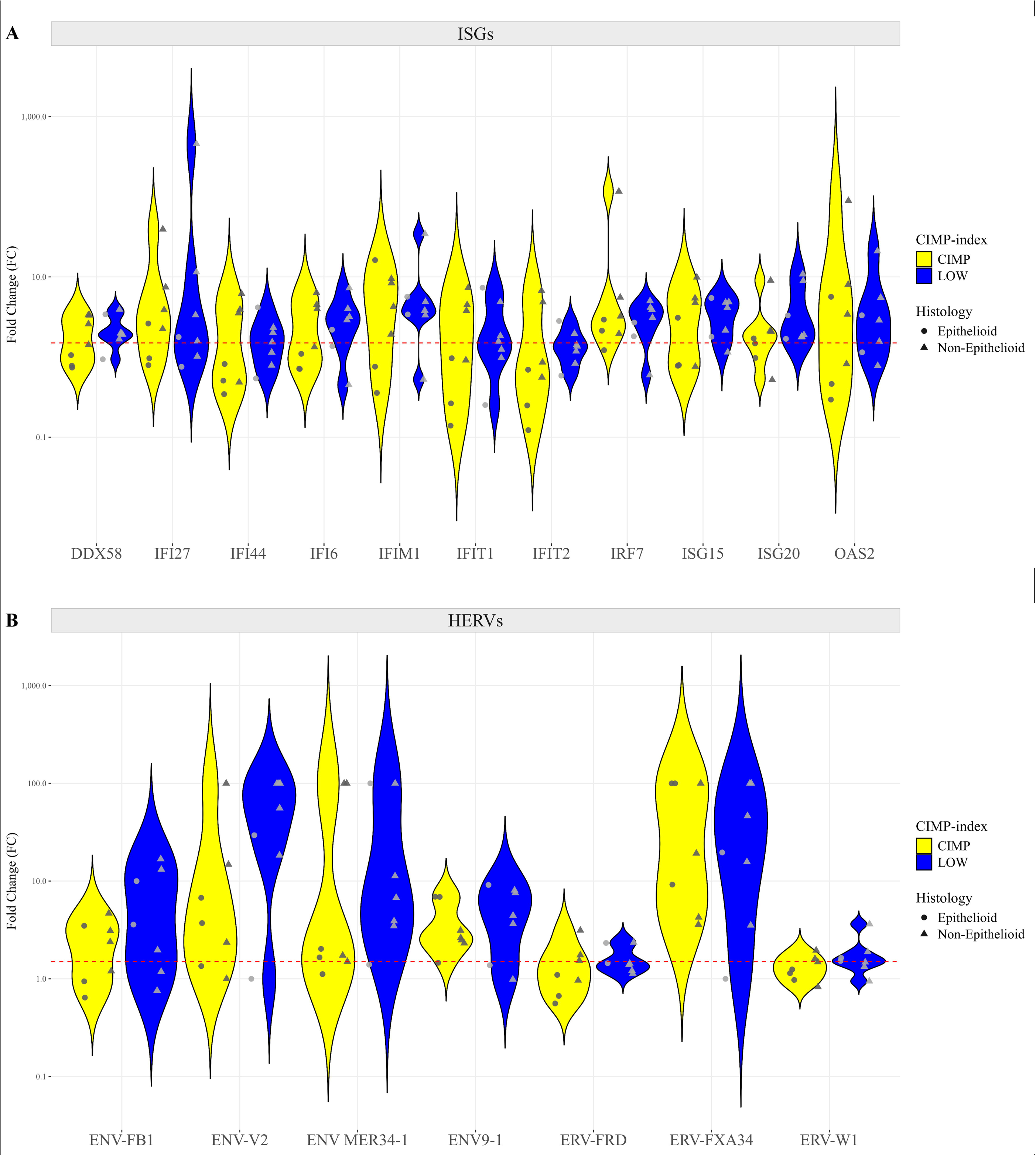
Modulation of viral mimicry-mediating genes expression in DHA-treated *vs* untreated CIMP and LOW PM cell lines. Starting from retrotranscribed total RNA of untreated and guadecitabine treated PM cell lines (#14), relative Real-Time PCR analyses were performed on (**A**) HERV (#7) and (**B**) ISG (#11) genes. Values are reported in the violin plots as FC of specific genes in treated *vs* untreated cell. Each symbol on the graph represents a cell line categorized by its CIMP-index: CIMP (yellow) and LOW (blue), and by its histopathological variant: E (circle) and non-E (triangle). Dashed line (red) represents a FC expression value ≥1.5.

### Integrative analysis of promoter methylation level and transcriptomic profiles in DHA treated *vs* untreated CIMP *and* LOW PM cell lines

To test whether the hyper-methylated immune-silenced profile associated with CIMP PM cell lines was reverted by DHA treatment, we performed integrative analyses of promoter methylation and transcriptomic profiling comparing DHA-treated *vs* untreated CIMP and LOW PM cell lines. This highlighted genes up regulated by promoter hypomethylation as the most impacted groups (**Figure 10 A**). Among the #467 hypo-methylated and up-regulated expressed genes in DHA-treated *vs* untreated CIMP cells, IPA core analysis revealed that treatment activated (Z-score ≥2) CP linked to the immune regulation functions, including the activation of antigen presentation machinery machinery (e.g., MHC class I/II antigen presentation), DC maturation, NK cell signaling, BCR signaling, cGAS-STING signaling pathway (**Additional File 5: Table S1)**. These results were further supported by the enrichment analysis of the #116 activated (Z-score ≥2) UR that identified the activation of immune-related BP along with those involved in the regulation of miRNA transcription, macromolecules metabolic processes and DNA transcription (**Additional File 5: Table S2, S3**). Conversely, in LOW PM cell lines, the intersection analysis of promoter methylation and gene expression, identified #49 genes (data not shown) as hypo-methylated and upregulated following DHA treatment.

**Figure 10.**
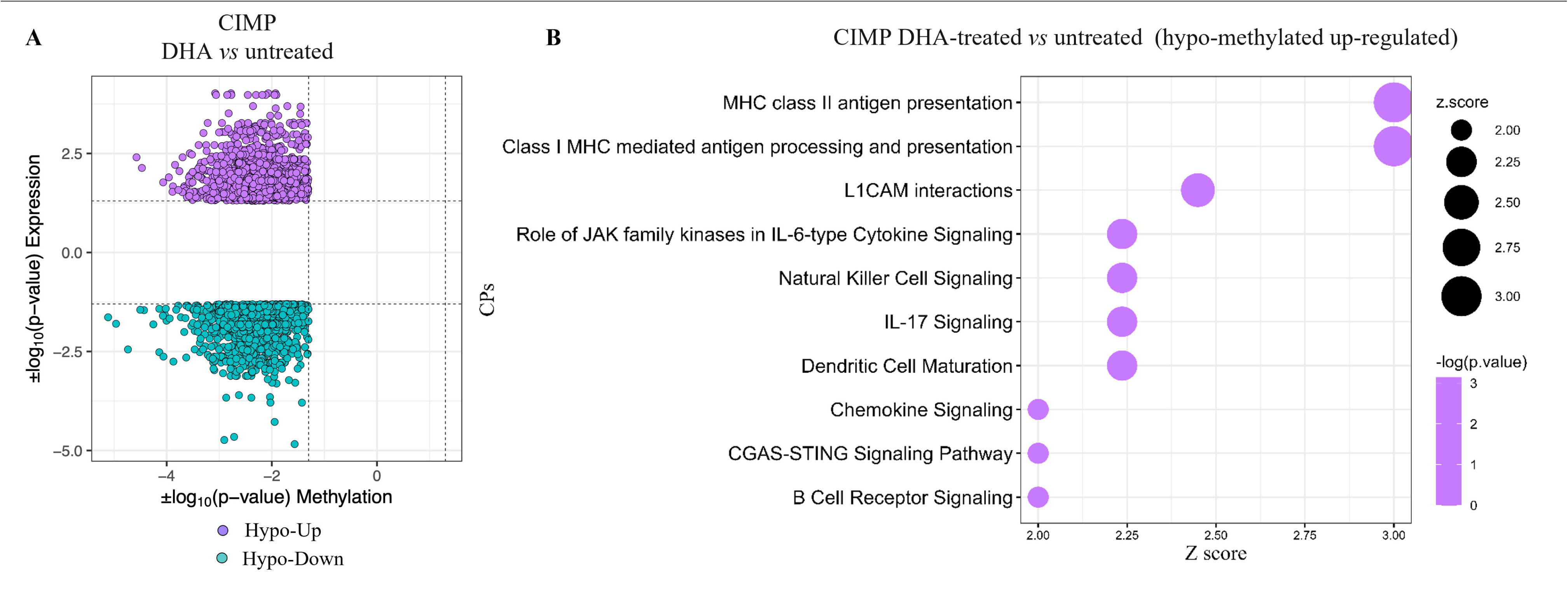
Integrative methylation and expression analysis. The starburst plot shows results of the integrative analysis of promoter methylation and transcriptomic data in DHA treated *vs* untreated CIMP PM cell lines. x and y axis represent the |log_10_(*p*-value)| of methylation and the |log_10_(*p*-value)| of expression (**A**). CP enriched by hypo-methylated and u*p-*regulated genes in DHA-treated *vs* untreated CIMP PM cell lines by IPA analysis (**B**).

## Discussion

In this study, we utilized a panel of cultured PM cells of different histology to define the contribution of tumor DNA methylation landscape in PM heterogeneity, and to provide preclinical evidence supporting the role of epigenetic remodeling to prospectively improve the efficacy of ICI therapy in PM patients.

To this end, we investigated methylation profiles of PM cell lines to stratify them according to their CIMP-index, in CIMP and LOW classes. Consistent with the notion that the CIMP index is independent from PM morphology (12), our stratification of PM cell lines did not depend upon their E or non-E histological subtype, but rather complements the histopathological classification of PM. Likewise, it was recently demonstrated that the histopathological classification only explains a fraction of the PM heterogeneity, since ploidy, adaptive immune response and CpG island methylation reflected the variations observed in the clinical behavior of PM patients (26).

Moreover, the integrated analysis of epigenetic and transcriptomic characterization of PM cell lines enabled us to explore which biological functions were associated with the two distinct DNA methylation-based classes. This analysis revealed that the CIMP profile included numerous hyper-methylated and silenced genes that impacted BP predominantly involved in shaping a more immune-compromised TME, compared to LOW PM cells.

Reinforcing the potential role of DNA methylation in the shaping tumor methylation profile and its immune contexture, are the results extrapolated from the TCGA database, that confirm the immune-favorable profile of LOW *vs* CIMP PM lesions. Along this line is the observation that low levels of tumor methylation were strongly associated with the presence of various immune cell subsets, including CD4^+^ regulatory T cells, and lymphocyte infiltration, across different cancer types such as breast cancer, head and neck tumors and lung adenocarcinoma (27). In line with these findings, Zou *et al*., demonstrated that the low CD8+ methylation TIL score (enriched CD8+ TILs) predicted better survival in colorectal cancer cohorts (28), reinforcing the concept that the methylation pattern may offer opportunities to model tumor immune compartment. In this context, the dysregulated methylation level can influence the immune function, and increased methylation may act as a biomarker to detect the response level to immunotherapy (29).

Based on these findings and considering the identified association between DNA hypermethylation and the immunosuppressive phenotype of CIMP PM cell lines, we investigated the remodeling activities of guadecitabine on immune-related patterns of CIMP and LOW PM cell lines. Interestingly, treatment with DHA both randomly reverted the methylation-driven immune-compromised profile of CIMP cell lines and reinforced the constitutive immune-favorable profile of LOW PM cell lines. Notably, among the BP commonly affected by DHA treatment in both CIMP and LOW PM cells was the cGAS-STING signaling pathway. This pathway has recently been proposed as a strategy to achieve stronger and more durable efficacy of ICI-based immunotherapy due to its ability to promote the release of type I IFN and multiple inflammatory cytokines (30). In line with these results are the modulation of HERVs and ISGs genes, considered potent inducers of the IFN-mediated viral mimicry response in PM (31), observed in both CIMP and LOW PM cell lines. These findings are consistent with the recent work of Sun and colleagues showing that DHA therapy can unleash viral mimicry in PM, which is furtherly promising since basal activation of this phenomenon is associated with better survival and clinical outcome in PM patients (32). In addition, guadecitabine enhances the expression of various genes involved in the antigen presentation machinery in both CIMP and LOW PM cell lines. This enhancement is crucial for reinvigorating anti-tumor CD8 T cells, restoring immune control of tumors, and improving immunotherapy efficacy in PM patients (33). Furthermore, guadecitabine promotes dendritic cell maturation, known for their potent antigen-presenting capacity (34), and NK cell signaling, which is being studied for its anti-tumor activity as an immunotherapeutic approach for PM. Beyond the reactivation of most IR-functions, influenced by hypomethylation of specific gene promoter regions, it is fair to mention the inhibitory action of guadecitabine on the pro-metastatic WNT/β-catenin signaling. This pathway is deeply implicated in PM pathogenesis (35) and correlated with immune exclusion in several human cancers (36). Additionally, guadecitabine negatively affected the major downstream effectors of the Hippo pathway, on the YAP/TAZ-axis, known to confer a proliferation advantage on PM cells via transcriptional regulation of cell cycle-related genes, as well as on tumorigenesis, progression, metastasis, and recurrence (37). Comprehensively, as already observed in previous *in vitro* studies (14), immune-modulation emerged as the preponderant beneficial effect of guadecitabine in PM cell lines, particularly in the immune-compromised CIMP ones.

## Conclusions

Our present study supports the role of DNA methylation in shaping the heterogeneous immune profile of PM cells, regardless of their histological classification. Further confirmatory analyses are granted in the clinical setting of PM; however, the pharmacologic immune remodeling induced by DHA in both PM methylation classes lays the foundations for the use of DHA in prospective clinical trials of precision epigenetic therapy.

## Additional files

**Additional Figure 1. Distribution of PM cell lines based on differentially methylated probes.** Dimensionality reduction was performed applying PCA on the DM probes among all PM cell lines. Each symbol on the graph represented a cell line categorized by its methylation status: CIMP (yellow) and LOW (blue) and by its histopathological variant: E-PM (circle) and non-E-PM (triangle).

**Additional Figure 2. Distribution of DNA methylation level in relation to CpG island regions and genomic regions in CIMP and LOW PM cell lines.** Bar graphs represented the distribution of DM probes across different CpG island regions (**A**, **B**) and functional regions including first exon (1stExon), 3’ untranslated region (3’UTR), 5’ untranslated region (5’ UTR), gene body (Body), exon boundaries (ExonBnd), 1500 bases upstream of the transcription site (TSS1500) and 200 bases upstream of the transcription site (TSS200) (**C**, **D**).

## List of abbreviations

BCR: B cell receptor
BP: biological processes
cGAS-STING: cyclic GMP-AMP synthase (cGAS)-stimulator of interferon genes (STING)
CIMP: CpG island methylator phenotype
CP: canonical pathways
CTLA-4: cytotoxic T lymphocyte antigen-4
DC: dendritic cell
DHA: DNA hypomethylating agent
DM: differentially methylated
E: epithelioid
E-score: epithelioid score
HERV: human endogenous retrovirus
ICI: immune-checkpoint inhibitors
ICOS: inducible Co-Stimulator
IPA: ingenuity pathway analysis
IR: immune-related
ISG: interferon stimulating gene
mAb: monoclonal antibody
MDSC: myeloid derived suppressor cells
MHC: major histocompatibility complex
NES: normalized enrichment scores
NK: natural killer
non-E: non-epithelioid
OS: overall survival
PCA: principal component analysis
PD-1: programmed cell death
PM: pleural mesothelioma **S-score** sarcomatoid-score
TCR: T cell receptor
TGF-β: transforming growth factor beta
Th17: T-helper (Th)-17
Th2: T-helper (Th)-2
TIL: tumor-infiltrating lymphocyte
TLS: tertiary lymphoid structure
TME: tumour microenvironment
UR: upstream regulators

## Declarations

### Ethics approval and consent to participate

Not applicable.

### Consent for publication

Not applicable.

### Availability of data and materials

All data analysed during this study are included in this published article as supplementary information files and/or are available from the corresponding author on reasonable request.

## Authors’ contributions

Conceptualization: M.F.L., A.C., S.C., M.M, A.M.D.G; Analysis: M.F.L., R.T., E.B., L.S., F.P., F.M., F.C.; Interpretation of data: M.F.L., A.C., S.C., R.T., F.P.C., T.M.R.N., R.M.; L.C. Writing: M.F.L., A.C., S.C.; Supervision: A.C., M.M., M.C., A.A. All authors have approved the submitted version.

## Acknowledgements

Not applicable.

## References

1. Zhou JG, Zhong H, Zhang J, Jin SH, Roudi R, Ma H. Development and Validation of a Prognostic Signature for Malignant Pleural Mesothelioma. Front Oncol. 2019;9:78.

2. Chiaro J, Antignani G, Feola S, Feodoroff M, Martins B, Cojoc H, et al. Development of mesothelioma-specific oncolytic immunotherapy enabled by immunopeptidomics of murine and human mesothelioma tumors. Nat Commun. 2023;14(1):7056.

3. Calabrò L, Morra A, Fonsatti E, Cutaia O, Amato G, Giannarelli D, et al. Tremelimumab for patients with chemotherapy-resistant advanced malignant mesothelioma: an open-label, single-arm, phase 2 trial. The Lancet Oncology. 2013;14(11):1104–11.

4. Calabrò L, Morra A, Fonsatti E, Cutaia O, Fazio C, Annesi D, et al. Efficacy and safety of an intensified schedule of tremelimumab for chemotherapy-resistant malignant mesothelioma: an open-label, single-arm, phase 2 study. The Lancet Respiratory Medicine. 2015;3(4):301–9.

5. Maio M, Scherpereel A, Calabrò L, Aerts J, Perez SC, Bearz A, et al. Tremelimumab as second-line or third-line treatment in relapsed malignant mesothelioma (DETERMINE): a multicentre, international, randomised, double-blind, placebo-controlled phase 2b trial. The Lancet Oncology. 2017;18(9):1261– 73.

6. Baas P, Scherpereel A, Nowak AK, Fujimoto N, Peters S, Tsao AS, et al. First-line nivolumab plus ipilimumab in unresectable malignant pleural mesothelioma (CheckMate 743): a multicentre, randomised, open-label, phase 3 trial. The Lancet. 2021;397(10272):375–86.

7. Peters S, Scherpereel A, Cornelissen R, Oulkhouir Y, Greillier L, Kaplan MA, et al. First-line nivolumab plus ipilimumab versus chemotherapy in patients with unresectable malignant pleural mesothelioma: 3-year outcomes from CheckMate 743. Annals of Oncology. 2022;33(5):488–99.

8. Minnema-Luiting J, Vroman H, Aerts J, Cornelissen R. Heterogeneity in Immune Cell Content in Malignant Pleural Mesothelioma. Int J Mol Sci. 2018;19(4):1041.

9. Santiago-Sánchez GS, Fabian KP, Hodge JW. A landscape of checkpoint blockade resistance in cancer: underlying mechanisms and current strategies to overcome resistance. Cancer Biology & Therapy. 2024;25(1):2308097.

10. Alay A, Cordero D, Hijazo-Pechero S, Aliagas E, Lopez-Doriga A, Marín R, et al. Integrative transcriptome analysis of malignant pleural mesothelioma reveals a clinically relevant immune-based classification. J Immunother Cancer. 2021;9(2):e001601.

11. Mannarino L, Paracchini L, Pezzuto F, Olteanu GE, Moracci L, Vedovelli L, et al. Epithelioid Pleural Mesothelioma Is Characterized by Tertiary Lymphoid Structures in Long Survivors: Results from the MATCH Study. IJMS. 2022;23(10):5786.

12. Blum Y, Meiller C, Quetel L, Elarouci N, Ayadi M, Tashtanbaeva D, et al. Dissecting heterogeneity in malignant pleural mesothelioma through histo-molecular gradients for clinical applications. Nat Commun. 2019;10(1):1333.

13. Maio M, Covre A, Fratta E, Di Giacomo AM, Taverna P, Natali PG, et al. Molecular Pathways: At the Crossroads of Cancer Epigenetics and Immunotherapy. Clinical Cancer Research. 2015;21(18):4040–7.

14. Lofiego MF, Cannito S, Fazio C, Piazzini F, Cutaia O, Solmonese L, et al. Epigenetic Immune Remodeling of Mesothelioma Cells: A New Strategy to Improve the Efficacy of Immunotherapy. Epigenomes. 2021;5(4):27.

15. Anichini A, Molla A, Nicolini G, Perotti VE, Sgambelluri F, Covre A, et al. Landscape of immune-related signatures induced by targeting of different epigenetic regulators in melanoma: implications for immunotherapy. J Exp Clin Cancer Res. 2022;41(1):325.

16. Lofiego MF, Piazzini F, Caruso FP, Marzani F, Solmonese L, Bello E, et al. Epigenetic remodeling to improve the efficacy of immunotherapy in human glioblastoma: pre-clinical evidence for development of new immunotherapy approaches. J Transl Med. 2024;22(1):223.

17. Noviello TMR, Di Giacomo AM, Caruso FP, Covre A, Mortarini R, Scala G, et al. Guadecitabine plus ipilimumab in unresectable melanoma: five-year follow-up and integrated multi-omic analysis in the phase 1b NIBIT-M4 trial. Nat Commun. 2023;14(1):5914.

18. Coral S, Covre A, Jmg Nicolay H, Parisi G, Rizzo A, Colizzi F, et al. Epigenetic remodelling of gene expression profiles of neoplastic and normal tissues: immunotherapeutic implications. Br J Cancer. 2012;107(7):1116–24.

19. Fortin JP, Labbe A, Lemire M, Zanke BW, Hudson TJ, Fertig EJ, et al. Functional normalization of 450k methylation array data improves replication in large cancer studies. Genome Biol. 2014;15(11):503.

20. Aryee MJ, Jaffe AE, Corrada-Bravo H, Ladd-Acosta C, Feinberg AP, Hansen KD, et al. Minfi: a flexible and comprehensive Bioconductor package for the analysis of Infinium DNA methylation microarrays. Bioinformatics. 2014;30(10):1363–9.

21. Ritchie ME, Phipson B, Wu D, Hu Y, Law CW, Shi W, et al. limma powers differential expression analyses for RNA-sequencing and microarray studies. Nucleic Acids Research. 2015;43(7):e47–e47.

22. Phipson B, Lee S, Majewski IJ, Alexander WS, Smyth GK. Robust hyperparameter estimation protects against hypervariable genes and improves power to detect differential expression. Ann Appl Stat. 2016;10(2).

23. Alam SkK, Astone M, Liu P, Hall SR, Coyle AM, Dankert EN, et al. DARPP-32 and t-DARPP promote non-small cell lung cancer growth through regulation of IKKα-dependent cell migration. Commun Biol. 2018;1(1):43.

24. Silva TC, Colaprico A, Olsen C, D’Angelo F, Bontempi G, Ceccarelli M, Noushmehr H. TCGA Workflow: Analyze cancer genomics and epigenomics data using Bioconductor packages. F1000Res. 2016; 29;5:1542.

25. Wu T, Hu E, Xu S, Chen M, Guo P, Dai Z, Feng T, Zhou L, Tang W, Zhan L, Fu X, Liu S, Bo X, Yu G. clusterProfiler 4.0: A universal enrichment tool for interpreting omics data. Innovation (Camb). 2021; 1;2(3):100141.

26. Becht E, Giraldo NA, Lacroix L, Buttard B, Elarouci N, Petitprez F, Selves J, Laurent-Puig P, Sautès-Fridman C, Fridman WH, de Reyniès A. Estimating the population abundance of tissue-infiltrating immune and stromal cell populations using gene expression. Genome Biol. 2016; 20;17(1):218.

24. Orozco Morales ML, Rinaldi CA, De Jong E, Lansley SM, Gummer JPA, Olasz B, et al. PPARα and PPARγ activation is associated with pleural mesothelioma invasion but therapeutic inhibition is ineffective. iScience. 2021;25(1):103571.

25. Kuleshov MV, Jones MR, Rouillard AD, Fernandez NF, Duan Q, Wang Z, et al. Enrichr: a comprehensive gene set enrichment analysis web server 2016 update. Nucleic Acids Res. 2016;44(W1):W90–7.

26. Mangiante L, Alcala N, Sexton-Oates A, Di Genova A, Gonzalez-Perez A, Khandekar A, et al. Multiomic analysis of malignant pleural mesothelioma identifies molecular axes and specialized tumor profiles driving intertumor heterogeneity. Nat Genet. 2023;55(4):607–18.

27. Qin Q, Zhou Y, Guo J, Chen Q, Tang W, Li Y, et al. Conserved methylation signatures associate with the tumor immune microenvironment and immunotherapy response. Genome Med. 2024;16(1):47.

28. Zou Q, Wang X, Ren D, Hu B, Tang G, Zhang Y, et al. DNA methylation-based signature of CD8+ tumor-infiltrating lymphocytes enables evaluation of immune response and prognosis in colorectal cancer. J Immunother Cancer. 2021;9(9):e002671.

29. Zhang C, Sheng Q, Zhao N, Huang S, Zhao Y. DNA hypomethylation mediates immune response in pan-cancer. Epigenetics. 2023;18(1):2192894.

30. Tian X, Xu F, Zhu Q, Feng Z, Dai W, Zhou Y, et al. Medicinal chemistry perspective on cGAS-STING signaling pathway with small molecule inhibitors. Eur J Med Chem. 2022;244:114791.

31. Chiappinelli KB, Strissel PL, Desrichard A, Li H, Henke C, Akman B, et al. Inhibiting DNA Methylation Causes an Interferon Response in Cancer via dsRNA Including Endogenous Retroviruses. Cell. 2016;164(5):1073.

32. Sun S, Qi W, Rehrauer H, Ronner M, Hariharan A, Wipplinger M, et al. Viral Mimicry Response Is Associated With Clinical Outcome in Pleural Mesothelioma. JTO Clin Res Rep. 2022;3(12):100430.

33. Lee HS, Jang HJ, Choi JM, Zhang J, de Rosen VL, Wheeler TM, et al. Comprehensive immunoproteogenomic analyses of malignant pleural mesothelioma. JCI Insight. 2018;3(7):e98575, 98575.

34. Cornelissen R, Lievense LA, Heuvers ME, Maat AP, Hendriks RW, Hoogsteden HC, et al. Dendritic Cell-Based Immunotherapy in Mesothelioma. Immunotherapy. 2012;4(10):1011–22.

35. Anani W, Bruggeman R, Zander DS. β-catenin expression in benign and malignant pleural disorders. Int J Clin Exp Pathol. 2011;4(8):742–7.

36. Luke JJ, Bao R, Sweis RF, Spranger S, Gajewski TF. WNT/β-catenin Pathway Activation Correlates with Immune Exclusion across Human Cancers. Clin Cancer Res. 2019;25(10):3074–83.

37. Zanconato F, Cordenonsi M, Piccolo S. YAP/TAZ at the Roots of Cancer. Cancer Cell. 2016;29(6):783– 803.

