## Supplementary figures and images for "DNA methylation status classifies pleural mesothelioma cells according to their immune profile: implication for precision epigenetic therapy"

### Additional Figure 1

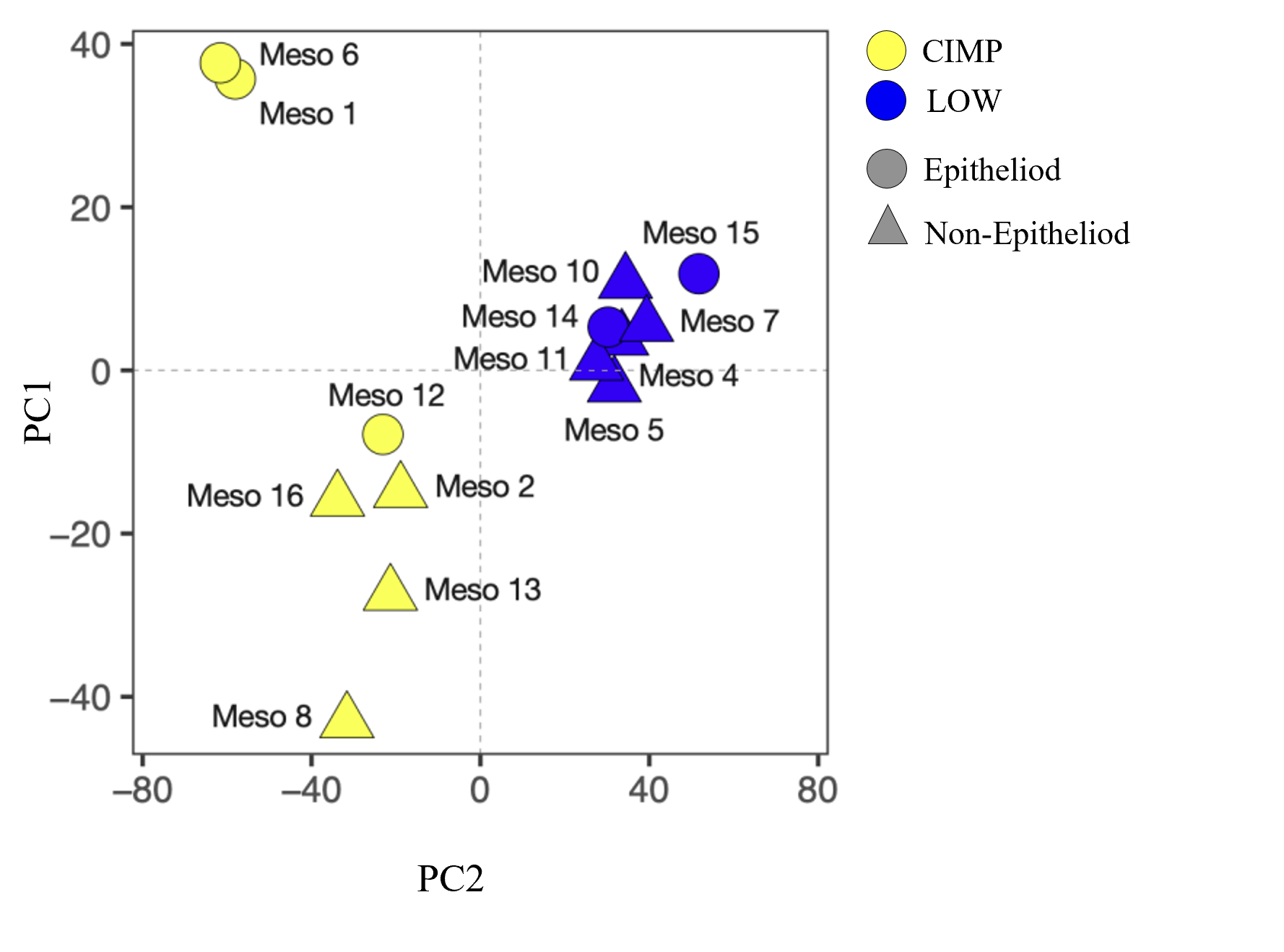

### Additional Figure 2

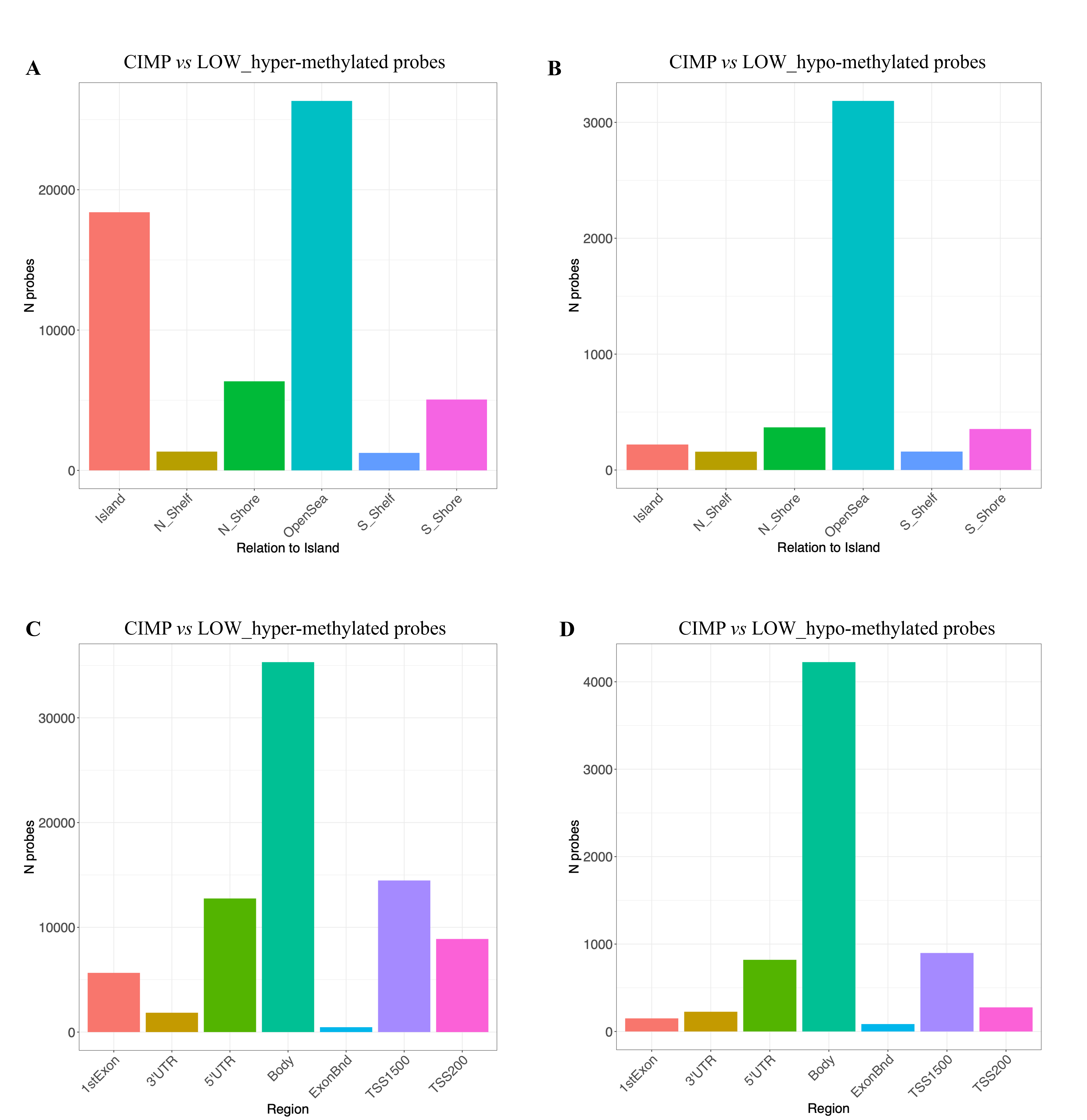
